# A disulfide constrains the ToxR periplasmic domain structure, altering its interactions with ToxS and bile-salts

**DOI:** 10.1101/2020.02.24.963330

**Authors:** Charles R Midgett, Rachel A Swindell, Maria Pellegrini, F Jon Kull

## Abstract

ToxR is a transmembrane transcription factor that, together with its integral membrane periplasmic binding partner ToxS, is conserved across the *Vibrio* family. In some pathogenic *Vibrios*, including *V. parahaemolyticus* and *V. cholerae*, ToxR is required for bile resistance and virulence, and ToxR is fully activated and protected from degradation by ToxS. ToxS achieves this in part by ensuring formation of an intra-chain disulfide bond in the C-terminal periplasmic domain of ToxR (dbToxRp). In this study, biochemical analysis showed dbToxRp to have a higher affinity for the ToxS periplasmic domain than the non-disulfide bonded conformation. Analysis of our dbToxRp crystal structure showed this is due to disulfide bond stabilization. Furthermore, dbToxRp is structurally homologous to the *V. parahaemolyticus* VtrA periplasmic domain. These results highlight the critical structural role of disulfide bond in ToxR and along with VtrA define a domain fold involved in environmental sensing conserved across the *Vibrio* family.

## Introduction

ToxR is the founding member of a group of transmembrane transcription factors (Schlundt *et al.*, 2017) and is conserved throughout the gram-negative *Vibrionaceae* family of marine bacteria (Reich and Schoolnik, 1994; Welch and Bartlett, 1998; Provenzano *et al.*, 2000; Wang *et al.*, 2002). Several *Vibrio* species are dependent on ToxR for bile resistance (Provenzano *et al.*, 2000), and a few pathogenic species have co-opted ToxR for virulence induction (Miller and Mekalanos, 1984; Hubbard *et al.*, 2016). *Vibrio parahaemolyticus* ToxR is important for colonization and virulence in mouse models (Hubbard *et al.*, 2016), where it induces type III secretion system expression through activating expression of VtrA and VtrB (Hubbard *et al.*, 2016). ToxS, an integral membrane periplasmic binding partner of ToxR, is also important for *V. parahaemolyticus* colonization in a mouse model (Hubbard *et al.*, 2016). In *V. cholerae*, ToxR is required for human colonization (Herrington *et al.*, 1988), most likely through induction of bile resistance (Provenzano *et al.*, 2000) and, in conjunction with TcpPH, virulence gene expression (Hase and Mekalanos, 1998).

ToxR regulates many genes in response to stimuli, including *ompT* and *ompU* (Bina *et al.*, 2003; Ante *et al.*, 2015). ToxR directly inhibits *ompT* expression (C C Li *et al.*, 2000) and activates *ompU* transcription (Crawford *et al.*, 1998), leading to a change in the outer-membrane porin composition (Miller and Mekalanos, 1988). ToxR is thought to activate virulence by augmenting TcpP activity, leading to *toxT* transcription (Hase and Mekalanos, 1998; Krukonis *et al.*, 2000; Krukonis and DiRita, 2003) and subsequent expression of cholera toxin and the toxin co-regulated pilus (Champion *et al.*, 1997; Krukonis *et al.*, 2000). Given the importance of ToxR and ToxS for both bile resistance and pathogenesis in the *Vibrio* family (Herrington *et al.*, 1988; Hubbard *et al.*, 2016), understanding how they interact with each other and respond to the environment is critical for understanding the disease processes and providing insight into how related proteins function in other pathogenic bacteria (Yang and Isberg, 1997; Peng Li *et al.*, 2016).

Previously, experiments in minimal media have shown *V. cholerae* ToxR activity can be regulated by the amino acid mixture of N, R, E and, S (NRES) (Mey *et al.*, 2012), or bile salts (Mey *et al.*, 2012; Midgett *et al.*, 2017). The NRES mixture led to an increase in ToxR and subsequent OmpU expression (Mey *et al.*, 2012; Mey *et al.*, 2015). In contrast, bile salts stimulated ToxR activity without increasing protein amounts (Mey *et al.*, 2015; Midgett *et al.*, 2017). This demonstrates ToxR activity can be regulated by two different mechanisms, one dependent on the Var/Csr system to increase ToxR mRNA and protein expression (Mey *et al.*, 2015), and a second to mobilize existing ToxR to become transcriptionally competent (Mey *et al.*, 2015; Midgett *et al.*, 2017). We hypothesize this potentially occurs via bile salts interacting with the ToxR periplasmic domain.

ToxS, an integral membrane protein with a periplasmic domain, augments ToxR activity (Miller *et al.*, 1989) and protects ToxR from degradation in conditions of stationary growth and alkaline pH (Almagro-Moreno *et al.*, 2015; Lembke *et al.*, 2018). Furthermore, ToxS is known to affect the formation of a disulfide bond in the ToxR periplasmic domain (Ottemann and Mekalanos, 1996; Fengler *et al.*, 2012). Investigations into the contribution of cysteines C236 and C293 on *V. cholerae* ToxR function showed single cysteine mutants were nonfunctional (Fan *et al.*, 2014), whereas a double mutant had decreased function in porin regulation, but no defects in virulence gene expression (Fengler *et al.*, 2012). Moreover, the double mutant was shown to be more susceptible to degradation than wild-type ToxR, even in the stabilizing presence of ToxS (Lembke *et al.*, 2018). In wild type ToxR, these cysteines are known to form either an intra-chain disulfide bond or a disulfide linked homodimer, which is present when *toxS* is deleted in *V. cholerae* (Ottemann and Mekalanos, 1995; Fengler *et al.*, 2012). Since *ΔtoxS* strains have defects in ToxR activity (Mey *et al.*, 2012; Fengler *et al.*, 2012; Midgett *et al.*, 2017), it follows that the intra-chain disulfide bonded, monomeric form of the ToxR periplasmic domain is the physiological active conformation.

While the intra-chain disulfide bonded conformation of ToxR is certainly the predominant form (Ottemann and Mekalanos, 1996; Fengler *et al.*, 2012), ToxR lacking a disulfide bond has been hypothesized to be present *in vivo* based on two prior studies. First, the periplasmic domain of ToxR and TcpP can form a disulfide linked heterodimer (Fan *et al.*, 2014), which can only occur if the two cysteines in ToxR are free to form a bond. Second, bile salts induce disulfide stress by inhibiting the activity of DsbA (Xue *et al.*, 2016), the chaperone responsible for inducing disulfide bond formation (Landeta *et al.*, 2018). Prior work in our laboratory showed the non-disulfide bonded ToxR periplasmic domain was destabilized by bile salts, however, the salts increased binding to the ToxS periplasmic domain (ToxSp). This led us to hypothesize the ToxR periplasmic domain has evolved to become destabilized by bile salts, leading to stronger ToxS binding and increased activity of ToxR (Midgett *et al.*, 2017).

To characterize and visualize the differences between the disulfide bonded and non-disulfide bonded forms of ToxR we expressed and purified the disulfide bonded conformer of the ToxR periplasmic domains (dbToxRp) from *V. cholerae* and *V. vulnificus* for biochemical and structural studies. The dbToxRp was able to bind ToxSp, and was destabilized by bile salts, although, unlike the non-disulfide bonded ToxR periplasmic domain (ToxRp), bile salts did not increase the binding of dbToxRp to ToxSp. The structure of dbToxRp showed the domain is globular composed of 2 α-helices, linked via the disulfide bond, and a β-sheet composed of 5 β-strands. Furthermore, we found that dbToxRp is structurally homologous to the periplasmic domain of VtrA (Peng Li *et al.*, 2016). A transmembrane transcription factor that shares similarities with ToxR as both have an OmpR DNA binding domain and a periplasmic domain that is involved in bile sensing (Kodama *et al.*, 2010; Peng Li *et al.*, 2016). Overall, these results demonstrate the important role of disulfide bond formation in interaction with ToxS, represent a first step in understanding the structure function relationship for virulence induction and bile resistance, and pave the way for a better understanding of the biochemical nature of activation of ToxR and related proteins.

## Results

### ToxR containing a disulfide bond is folded and active

The *V. cholerae* disulfide ToxR periplasmic domain (Vc-dbToxRp) construct was confirmed to be structurally and functionally similar to the previously analyzed ToxRp (Midgett *et al.*, 2017) by comparing NMR ^1^H,^15^N HSQC’s, by determining if CDC and cholate could lower the melting temperature, and assessing its ability to bind chitin binding domain-intein tag ToxS periplasmic domain (CBDI-ToxSp) (Fig. 1). A comparison of the Vc-dbToxRp HSQC with the previously collected HSQC of the ToxRp shows the two conformations share a common core fold (Fig. 1a), as demonstrated by a series of well dispersed peaks that overlap in the two conformations. About 10 extra intense peaks are observed for Vc-ToxRp in the central region of the spectrum, consistent with random coil shifts corresponding to the unfolding of the short helix α2. Differential scanning fluorometry in the presence of cholate and chenodeoxycholate (CDC) showed Vc-dbToxRp was destabilized by the bile salts, with CDC having the greatest effect (Fig. 1b), demonstrating the disulfide bonded conformer was destabilized in a manner similar to the non-disulfide bonded form as previously described (Midgett *et al.*, 2017). Finally, a pull down showed the Vc-dbToxRp interacted with the CBDI-ToxSp (Fig. 1c). Taken together, these tests confirm that Vc-dbToxRp is structurally and functionally similar to Vc-ToxRp.

**Figure 1:**
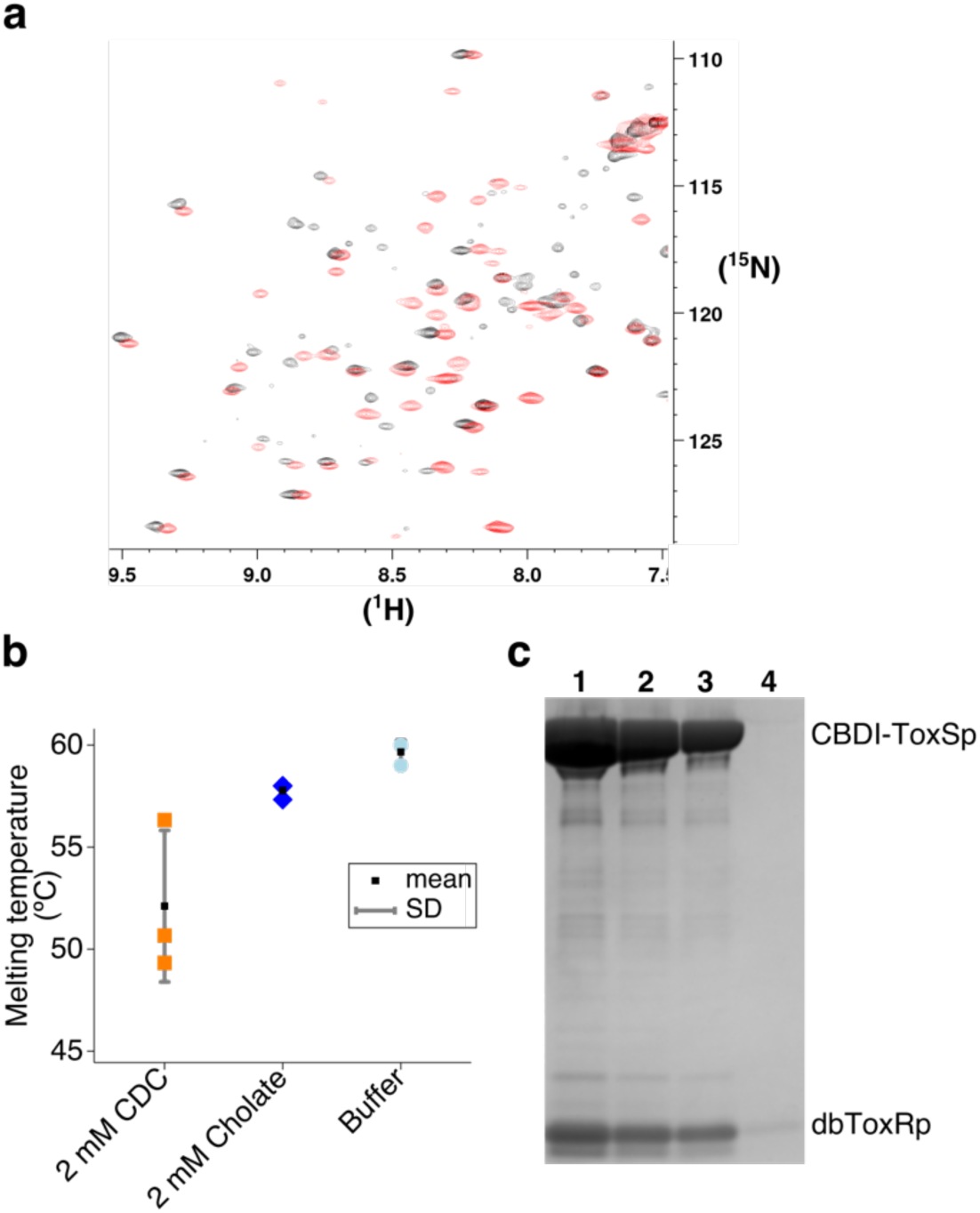
Validation of the dbToxRp construct showing it is similar in structure to ToxRp, destabilized by bile salts, and can bind ToxSp. **a.** A HSQC of 15N-dbToxRp in black overlaid with a HSQC of 15N-ToxRp in red. The NMR data was collected on a Bruker 700 MHz instrument. **b**. Melting temperatures of dbToxRp when treated with 2 mM cholate and chenodeoxycholate as measured by DSF; mean ± SD, n=3. **c.** A colloidal blue stained gel showing the dbToxRp can be pulled down by CBDI-ToxSp. Lane 1-3: serial dilutions 1x-1/4x of the CBDI-ToxSp pull down with dbToxRP, Lane 4: dbToxRp with beads only.

### dbToxRp shows increased binding to ToxSp, unaffected by bile salt

Previous work in our laboratory revealed, somewhat unexpectedly, that the bile salt CDC increased the interaction between the periplasmic domains of ToxR and ToxS (Midgett *et al.*, 2017). To test if CDC also increases the affinity of Vc-dbToxRp to CBDI-ToxSp we performed pull downs with ToxRp and Vc-dbToxRp in the presence and absence of CDC. Similar to previous results, CDC increased the amount of Vc-ToxRp pulled down by CBDI-ToxSp (fold change of 4.0 ± 2.0). However, CDC had a negligible effect on the amount of Vc-dbToxRp pulled down (fold change of 1.6 ± .5), indicating Vc-dbToxRp binding to ToxS is unaffected by bile salts (Fig. 2a). To compare the amounts of the ToxR periplasmic domains pulled down in the experiments we ran selected pull downs on the same gel to calculate relative amounts. Interestingly, the presence of a disulfide bond increased the interaction between Vc-dbToxRp and CBDI-ToxSp. The results clearly showed Vc-dbToxRp with CDC was pulled down more than Vc-ToxRp in in the presence or absence of CDC. As when the amount of Vc-dbToxRp pulled down was set to one, Vc-ToxRp was pulled down .22 ± .11 in comparison to Vc-dbToxRp, and Vc-ToxRp with CDC was pulled down .41 ± .05 relative to Vc-dbToxRp (Fig. 2b). These results suggest the presence of a disulfide bond in the ToxR periplasmic domain significantly increases the affinity of ToxR for ToxS.

**Figure 2:**
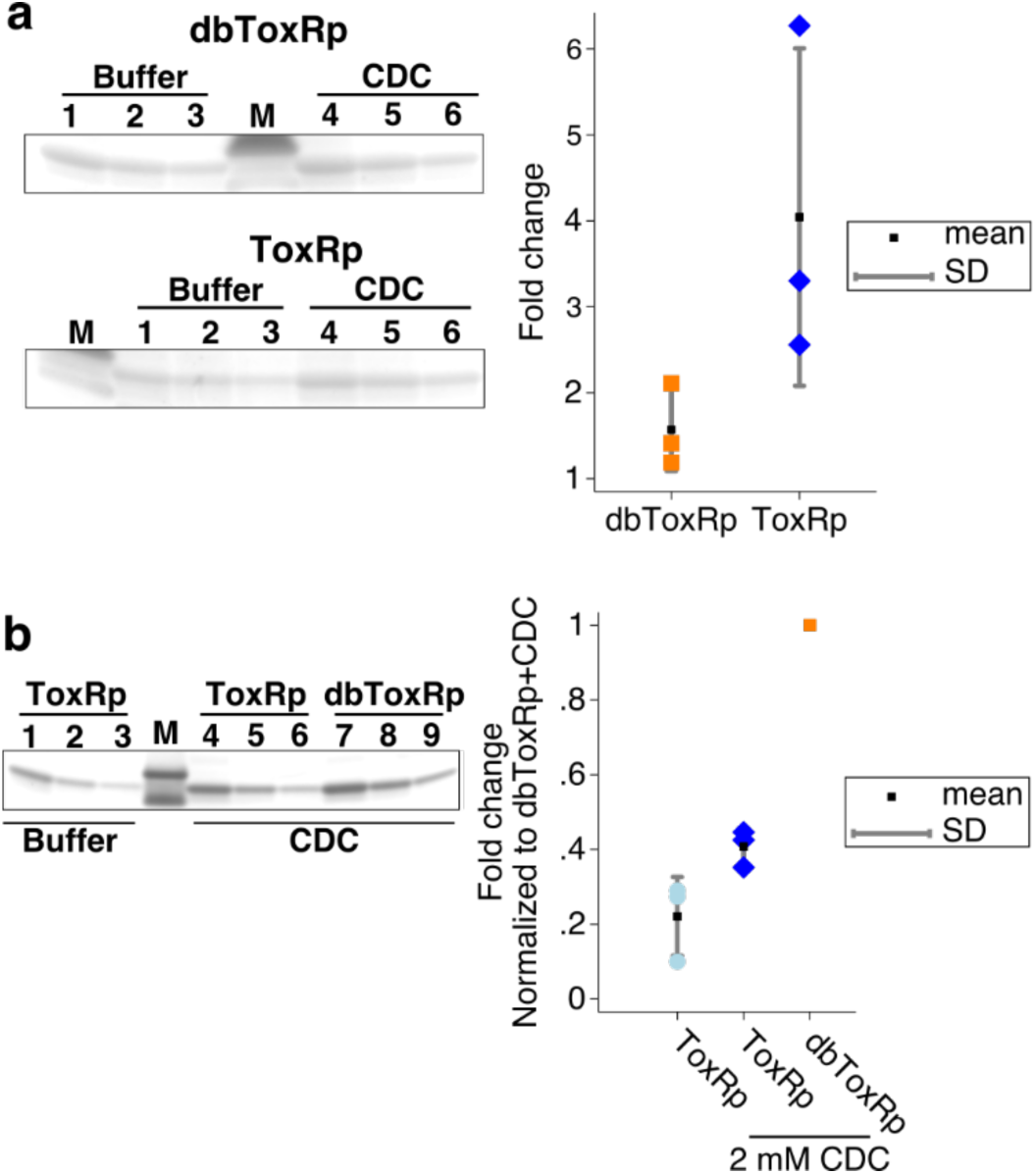
dbToxRp binds to ToxSp 2-4 fold more than ToxRp and the binding is not effected by bile salts. **a.** Left top panel a colloidal blue stained gel excerpt showing the changes in the amount of dbToxRp when the domain was treated with buffer versus 2 mM CDC. Left bottom panel a colloidal blue stained gel excerpt showing the changes in ToxRp pulled down when the domain was treated with buffer or 2 mM CDC. Lanes 1-3 serial dilutions of the CDC treated pull down and 4-6 serial dilutions of the pull down with buffer. M is the lane with the marker. Right panel is a graph showing the relative amount of the domains pulled down by the compounds versus buffer; mean ± SD, n=3. **b.** Left panel gel of the periplasmic domains pulled down from the above experiments to determine the relative amount of ToxRp pulled down relative to dbToxRp. Graph of the results normalized to the pull down performed with dbToxRp + 2 mM CDC (The intensity of dbToxRp 1x dilution band was set to one in each trial); mean ± SD, n=3.

### Crystal structures reveal the structural role of the ToxR periplasmic domain disulfide bond

The crystal structure of *V. vulnificus* intra-chain disulfide ToxR periplasmic domain (Vv-dbToxRp) was solved by molecular replacement using a SeMet labeled structure to 1.25 Å, revealing an α/β fold with a 5 stranded β-sheet with one side packing against two α-helices (Fig. 3a). The two helices are connected by a disulfide bond between Cys232 and Cys289. The structure is well ordered, and electron density for the disulfide bond is clearly visible in a composite omit map of the native structure (Fig 3b first panel), with only density of the α2-β5 loop being poorly defined (Fig 3b middle panel). The α2-β5 loop has the highest B-factors in the structure, indicating conformational flexibility (Fig 3b last panel). We hypothesize a major role of the disulfide bond is to stabilize the structure, and importantly the ToxS binding region, by constraining this flexible loop as well as the relatively short helix α2 which follows it.

**Figure 3:**
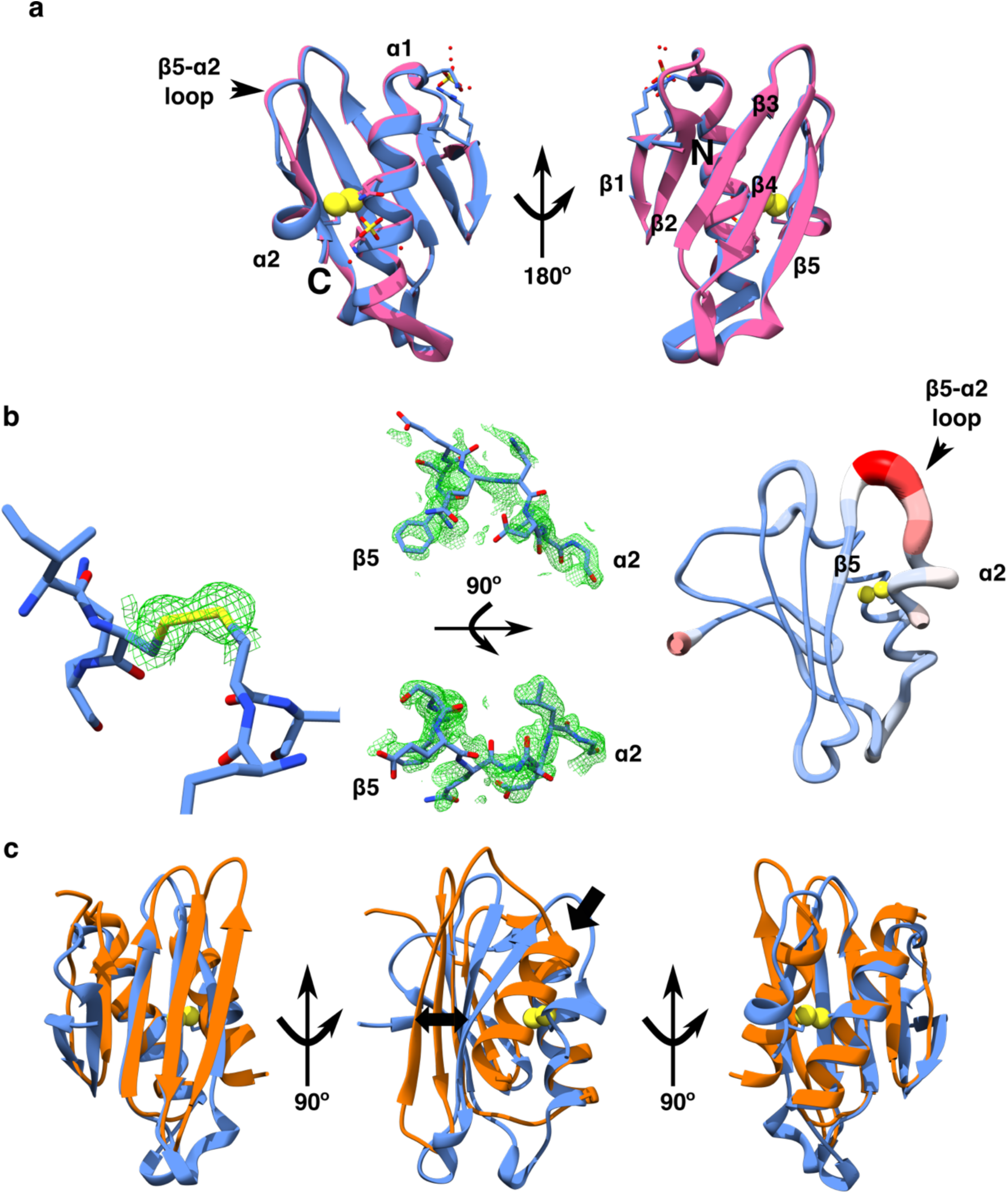
The structure of the *V. vulnificus* dbToxRp showing the electron density around the disulfide, the α2-β5 loop disorder, and structural homology with the VtrA periplasmic domain. **a.** Alignments of the SeMet-Vv-dbToxRp (pink) with the native Vv-dbToxRp (blue). The secondary structure elements are labeled along with the N- and C-termini. **b.** Left to right; the omit map density surrounding the disulfide bond displayed as green mesh. The density from the composite omit map around the loop displayed as a green mesh. Top panel is the side view and the bottom panel is the top view of the loop. Representation of the B-factors showing the β5-α2 loop has the highest B-factors in the structure. **c.** Overlay of the VtrA structure in orange and the Vv-dbToxRp structure in blue. A single arrow points to the last helix in VtrA, and a double arrow shows the displacement of the last strand between VtrA and ToxR structures.

### VtrA and ToxR periplasmic domains are structural and functional homologues

Using the Vv-dbToxRp structure, we ran a DALI search (Holm and Laakso, 2016) to look for structural homologues. Not surprisingly given the prevalence of α/β folds in proteins, DALI identified a number of functionally unrelated proteins with structural similarity. Interestingly, the periplasmic domain of VtrA from *V. parahaemolyticus* (Peng Li *et al.*, 2016) was ranked 118^th^ by the DALI server (Table S1). Despite having little sequence identity and an RMSD of 3.0-3.3 Å it is clearly a structural homolog (Fig. 3c). Not only do both domains have the same topology and overall structural motif, but both are C-terminal periplasmic domains of transmembrane transcription factors involved in environmental sensing (Welch and Bartlett, 1998; Mey *et al.*, 2012; Peng Li *et al.*, 2016; Midgett *et al.*, 2017). The main difference between the two structures is the position of the last β-strand and α-helix. In VtrA the last helix is longer, pushing the last β-strand further away from the core structure. Also, as the periplasmic domain of VtrA does not contain cysteines, there can be no stabilizing disulfide bond. Based on the clear structural and functional similarities, we propose the periplasmic domains of ToxR and VtrA represent a common domain utilized in sensing the extra-cellular environment.

### dbToxRp lacks a dimerization interface

The periplasmic domain of ToxR has been proposed to mediate dimerization, which is thought to be essential for activity. Prior studies replacing the periplasmic domain of ToxR with proteins known to dimerize, resulted in active constructs (DiRita and Mekalanos, 1991; Ottemann and Mekalanos, 1995; Kolmar *et al.*, 1995; Dziejman *et al.*, 1999). Therefore, it was somewhat surprising that the Vv-dbToxRp structure did not form a crystallographic dimer, nor did it illuminate a biologically relevant dimer interface. Close examination of the structure revealed two crystal contacts that could be potential dimerization interfaces (Fig. S1). The first involves the loop between β4 and β5, which makes a contact with the loop between β2 and α1 as well as the loop between β3 and β4 on a neighboring molecule. However, given that there are few hydrogen bonding or charge-charge interactions between the two protein molecules, and that dimerization in this area would position the N-termini on opposite sides of the dimer, an interface here would essentially sterically preclude both proteins being membrane anchored. Another possible interaction is between β1 and the β2-α1 loop on an adjacent monomer. While this would position both N-termini toward the membrane, the interaction is mediated by a sulfate ion and there are a few interactions between the two monomers. Analysis using PISA (Krissinel and Henrick, 2007) also corroborates that both of these interfaces are most likely biologically irrelevant. Therefore, we conclude the ToxR periplasmic domain is most likely monomeric and any dimerization is mediated by other factors, for example, via interaction with ToxS.

## Discussion

Here we have biochemically characterized and solved the structure of the ToxR periplasmic domain with an intra-chain disulfide bond. This conformation was found to be similar to the non-disulfide conformation as confirmed by NMR, indicating both constructs consist of a stable core fold. In addition, the disulfide conformer is destabilized by bile salts, similar to previously published results (Midgett *et al.*, 2017). However, in contrast to ToxRp (Midgett *et al.*, 2017), bile salts had no effect on the binding of the dbToxRp conformer to ToxSp. Furthermore, dbToxRp bound ToxSp about 2-4 fold more in our pull down assay than the non-disulfide bond conformer, suggesting the disulfide bond stabilizes the ToxS binding interface.

The dbToxRp structure is clearly homologous to the periplasmic domain of VtrA (Peng Li *et al.*, 2016). Furthermore, given VtrA and ToxR share functional similarities we can hypothesize how ToxR and ToxS interact based on what is known about the VtrA system. Previously, the VtrA periplasmic domain was crystallized with its binding partner the VtrC periplasmic domain. The structure showed the proteins interact in two ways, via parallel beta strand hydrogen bonds between the β5 strand of VtrA sheet and the most N-terminal β-strand of VtrC beta-barrel (arrow 1 in Fig 4a left panel), as well as β-sandwich like interaction between the VtrA β-sheet and the C-termnal region of the VtrC β-barrel (arrow two in Fig 4a left panel) (Peng Li *et al.*, 2016). Based on this, we predict ToxS is a structural homolog of VtrC and the ToxR periplasmic domain will bind ToxS in a similar manner (Fig. 4a right panel). Despite these structural similarities there are differences between these two regulatory pairs. First, VtrA requires VtrC for expression (Peng Li *et al.*, 2016) whereas ToxR can be expressed by itself (Ottemann and Mekalanos, 1996; Mey *et al.*, 2012; Fengler *et al.*, 2012; Midgett *et al.*, 2017). Second, ToxR depends on the formation of an intra-chain disulfide bond to properly fold while the VtrA periplasmic domain is cysteine free. It would be interesting to determine if the ToxR periplasmic domain could be stabilized without a disulfide bond and what effect that would have on ToxR function.

**Figure 4:**
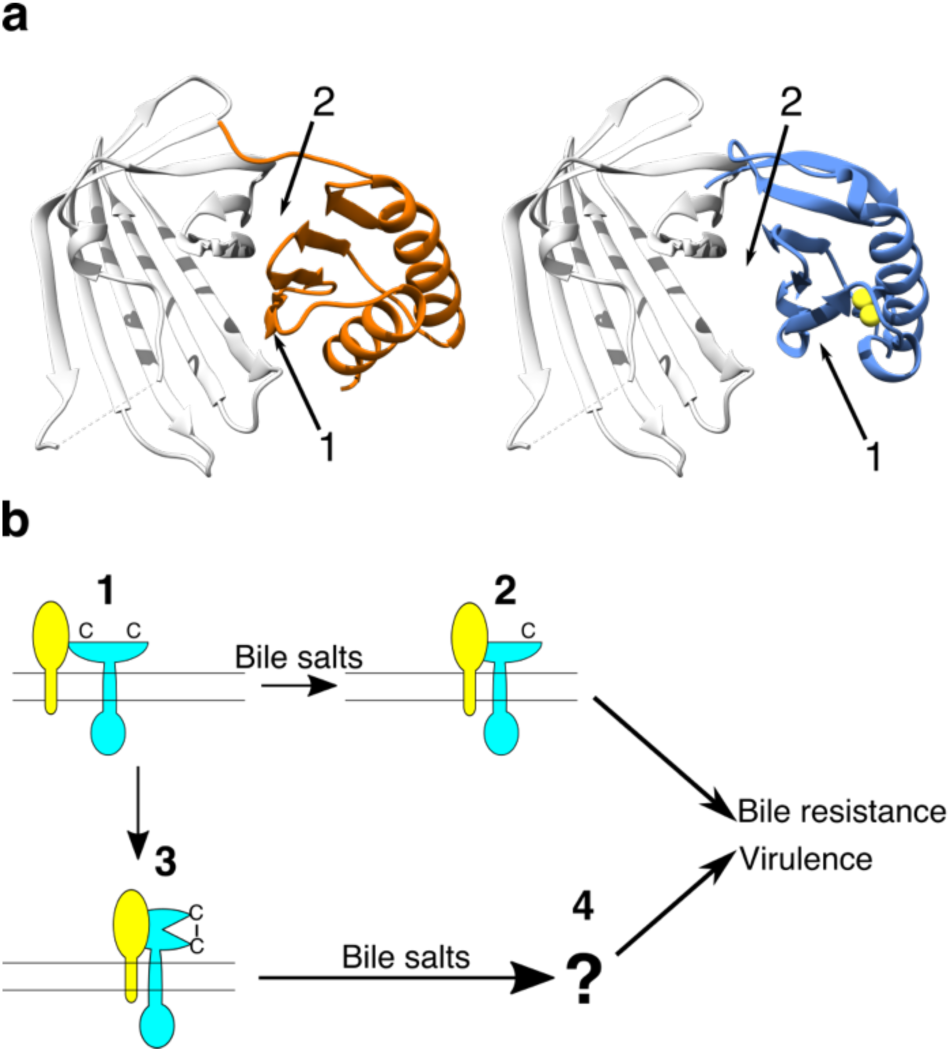
Models of potential interactions between ToxR and ToxS periplasmic domain using the VtrAC structure, and how ToxR and ToxS interact to induce virulence as well as bile resistance. **a.** The left panel shows the VtrAC (VtrA orange, VtrC in gray) structure with arrows pointing to the interfaces between the proteins. Arrow 1 points to the VtrA β5-strand that extends the VtrC β-sheet, and arrow 2 points to VtrA β-sandwich interaction with VtrC. In the right panel the Vv-dbToxRp structure (blue) was aligned with VtrA in the VtrAC structure, VtrC is depicted in gray. The numbered arrows point to the β5 strand (1) and the potential β-sheet interface (2). **b.** A model of the potential interactions between ToxR, ToxS, and bile salts. The non-disulfide bonded form of ToxR interacts weakly with ToxS (1). However, bile salts increase the interaction of the non-disulfide bonded ToxR to ToxS leading to ToxR activation (2). In addition, ToxS favors the formation of the intra-chain periplasmic disulfide bond conformation increasing the affinity between ToxR and ToxS (3). The complex of the periplasmic disulfide bonded ToxR with ToxS in the presence of bile salts increases ToxR activity by an unknown mechanism (4).

By combining our results with insight gleaned from the VtrAC structure, we can begin to explain how ToxR and ToxS might interact. At least in vitro, ToxR can exist in three forms, a monomer with an intra-chain disulfide bond, a disulfide-free monomer, and a (likely non-physiological) disulfide bonded homodimer. Disulfide formation in the ToxR monomer apparently constrains the β5-α2 loop and the adjacent α2 helix. This, based on structural homology with VtrAC, stabilizes the binding interface between ToxR and ToxS. Such an increase in stability is consistent with observations regarding the double cysteine mutant degradation in Lembke et al. (Lembke *et al.*, 2018). Interestingly it has been observed that ToxS enhances ToxR intra-chain disulfide bond formation (Ottemann and Mekalanos, 1996). How this occurs in the context of the above results is unexplored.

The structure of the dbToxRp provides a potential explanation to why breaking the disulfide bond interferes with ToxS binding and how bile salts can modulate the interaction. In the absence of intra-chain disulfide bond formation, ToxR is destabilized and it is unlikely the short α2 helix would form. Additionally, α2 might assume an alternate conformation in which it blocks interaction with ToxS. Given bile salts increase the ToxRp-ToxSp interaction suggests bile salts disrupt whatever alternate structure α2 forms in the absence of the disulfide bond, thereby allowing ToxS binding. In dbToxRp, the loop and α2 are already in a conformation that stabilizes the interface, rather than blocking it, and bile salts have no effect on ToxS binding.

This leaves open the question of how ToxS augments ToxR activity. ToxR is thought to be active as a dimer and its periplasmic domain has been presumed to mediate dimerization (DiRita and Mekalanos, 1991; Ottemann and Mekalanos, 1995; Kolmar *et al.*, 1995; Dziejman *et al.*, 1999). However, our structural analysis failed to find a dimer interface, arguing for other determinants of dimerization. ToxS is the most likely candidate for this, and we propose that disulfide bond dependent interaction of ToxR with ToxS results in formation of a dimer of ToxR and ToxS dimers, forming a complex competent for ToxR activation. Future structural studies aimed at determining the structures of ToxSp as well as that of the complex between dbToxRp and ToxSp should clarify if this is the mechanism for dimerization.

Alternately, given ToxR binds DNA, DNA could very well provide the impetus for ToxR oligomerization. In such a model ToxR binds DNA, and weak protein-protein interactions could promote dimerization or higher order oligomers, as seen in experiments with VtrA (Okada *et al.*, 2017). Yet another possibility is DNA organizes ToxR in the membrane without specific interactions between domains. Determining which might be the case will require reconstruction of full-length ToxR in a membrane and in the presence of DNA.

In summary, our results suggest a model in which ToxR is a monomer in the membrane. The non-disulfide bonded conformation has a low affinity for ToxS, and the interaction between the two proteins increases in the presence of bile salts, leading to increased ToxR activity. When ToxR contains an intra-chain disulfide bond, its affinity for ToxS is increased and unaffected by bile salts, leaving open the question of how bile salts activate ToxR (Fig. 4b).

## Materials and Methods

### Cloning of ToxR periplasmic domains

The sequences of the ToxR periplasmic domains from *V. vulnificus, V. parahaemolyticus, V. fischeri, V. harveyi*, and *Photobacterium profundum* were identified by using the ToxR periplasmic sequence from *V. cholerae* (T199-E294). The six ToxR periplasmic domains were codon optimized and synthesized, from 5’ to 3’, with a NcoI site, a 6xHis N-terminal tag followed by the coding sequence, a BamHI site, all flanked by primer sites to amplify the constructs. The constructs were PCR amplified, cut with the appropriate restriction enzymes, and the digested constructs were purified using a PCR clean up kit (Qiagen). The pET16b plasmid was digested with the same restriction enzymes, treated with CIP (NEB), and purified using a PCR clean up kit (Qiagen). The inserts were ligated into the plasmid using the Quick Ligase (NEB) and the reaction was used to transform DH5α’s. Colonies from the transformation were subjected to colony PCR to determine if the plasmids contained an insert of the appropriate size. Then selected colonies were cultured overnight for mini-preps following manufacture instructions (Qiagen). The resulting plasmids were sequence verified using a primer for the T7 promoter.

### Expression and purification

The plasmids with the different ToxR periplasmic domains were transformed into Shuffle T7 Express cells. The strains were double selected to produce stable expression strains (Sivashanmugam *et al.*, 2009). After double selection the strains were checked for production of soluble protein, by performing a scaled down purification. The ToxR periplasmic domain from *V. vulnificus* was found to produce the greatest amount of soluble protein. Production cultures of *V. vulnificus* and *V. cholerae* ToxR periplasmic domains were started by picking a colony from a freshly streaked plate incubated overnight at 37 °C, and inoculating 2ml of ZYP-0.8G media (Studier, 2005) with 200 *µ*g/ml of carbenicillin, and incubated overnight at 30 °C. The next morning the culture was used to inoculate Terrific Broth Modified (Fisher Scientific) media supplemented with 2 mM MgSO4 and 200 *µ*g/ml carbenicillin at 1:250, then grown to and OD600 of 1-1.2. The cultures were centrifuged at 600 xg for 10 minutes, at 25 °C, with the brake turned off. The cells were resuspended in an equal volume of M9 media with 100 *µ*g/ml carbenicillin and grown for 1 h at 37 °C. The cultures were induced with 1 mM IPTG and incubated overnight at 25 °C with loose covers to allow gas exchange.

For SeMet labeling the frozen culture was restreaked onto a plate containing 200 *µ*g/ml of carbenicillin and incubated overnight at 37 °C. In the morning a starter culture from a single colony in ZYP-0.8G media with 200 *µ*g/ml of carbenicillin incubated at 37 °C. In the evening the culture was used to inoculate cultures at 1:12.5 ratio of M9 media with 100 *µ*g/ml carbenicillin. The culture was grown overnight at 30 °C. The next morning the culture was used to inoculate M9 media at a ratio of 1:20. The culture was incubated at 37 °C till an OD600 of .5. Then 25 mg/L of lysine, phenylalanine, and threonine; 12.5 mg/L of isoleucine, leucine, and valine; and 15 mg/L of seleno-methionine were added to the culture. The culture was grown for 15 minutes then induced with 1 mM IPTG and incubated at 25 °C with a loose cover allowing air flow overnight.

For ^15^N and ^13^C labeling the cultures were started from a single fresh colony in ZYP-0.8G media with 200 *µ*g/ml of carbenicillin overnight at 30 °C. The cultures were used to inoculate 1:250 TB media and incubated at 37 °C till an OD600 of 2. The cultures were centrifuged at 600 xg at 25 °C for 20 minutes with the brake turned off. The supernatant was discarded and the cells were resuspended in an equal volume of M9 media with 100 *µ*g/ml carbenicillin. The resuspended cells were added to M9 media with 100 *µ*g/ml carbenicillin, 3 g/L 15N-NH4Cl, and 10 g/L of U-13C-glucose. The cultures were incubated at 37 °C for 1 h then induced with 1 mM IPTG and incubated at 25 °C overnight.

To purify the protein the cells were harvested by centrifugation at 4500 xg, at 4 °C. The cells were resuspended in lysis buffer (wash buffer (20 mM HEPES pH 7.4, 20 mM imidazole, 200 mM NaCl) with a Complete tablet EDTA free (Roche), 3 mM MgSO4, 1.5 mM EDTA). The cells were lysed with three passes through a French press. The lysate was clarified by centrifugation at 100,000 xg for 45 minutes. The resulting supernatant was filtered using a .45 *µ*m filter before purification. The protein was captured using a His-Trap column (GE Healthcare). The column was washed with 10 CV’s of wash buffer, 10 CV’s of high salt wash buffer (20 mM HEPES pH7.4, 20 mM imidazole, 1 M NaCl), 10 CV’s of wash buffer, followed by 9 CV’s of 9% elution buffer (20 mM HEPES pH 7.4, 500 mM imidazole, 200 mM NaCl). The protein was eluted using a gradient to 100% elution buffer over 4 CV’s with a 2 CV hold at 100%. The relevant fractions were pooled and flowed over an equilibrated thio-propyl sepharose 6B (GE Healthcare) column in order to bind protein with free cysteines. The flow through was collected and concentrated to about 2 ml. The final purification was performed using a S75 16/600 (GE Healthcare) column with the gel filtration buffer (20 mM HEPES pH 7.4, 200 mM NaCl for crystallization and biochemistry, or 20 mM KPO4 pH 7.4, 200 mM NaCl for NMR). The relevant fractions were pooled and concentrated as required.

To obtain the non-disulfide bonded form of the ToxR periplasmic domain the thio-propyl column was washed with 2 CV’s of wash buffer, then with 3 CV’s of wash buffer with 100 mM DTT. Two more CV’s of wash buffer with DTT were added to the column and incubated overnight at 16 °C. The following morning the protein was eluted and concentrated to about 2 ml for gel filtration as above.

### Differential Scanning Fluorometry

Differential scanning fluorometry was performed to assess the stability of the *V. cholerae* dbToxRp in the presence and absence of additives as described (Midgett *et al.*, 2017). STATA15 was used to analyze the data. The results are reported from three independent experiments as mean ± standard deviation.

### Pull downs

Pull downs were performed as described (Midgett *et al.*, 2017). Briefly, the chitin binding domain-intein tagged ToxS periplasmic domain (CBDI-ToxSp) was captured on chitin beads from a lysate. After washing the beads purified ToxR periplasmic domain with and without the disulfide bond were added to the tubes with and without 2 mM sodium chenodeoxycholate. After the final wash SDS sample buffer was added to the beads and the tubes were boiled. Samples were diluted and run on a gel with the results quantified and statistics determined using STATA15. The results are from three independent experiments.

### Nuclear Magnetic Resonance

^1^H,^15^N HSQ(Schleucher *et al.*, 1994) were acquired at 298 K, on a 700 MHz Avance NMR spectrometer equipped with a 5 mm TCI cryoprobe, utilizing 16 scans and 2048 x 256 points. The ToxR periplasmic domain without the disulfide bond was analyzed at a concentration of 580 *µ*M in 20 mM phosphate buffer pH 6.9, 200 mM NaCl, .2 mM TCEP, .02% NaN_3_, .1 mM Pefablock and 1.7% D_2_O. The disulfide bonded ToxR periplasmic domain was analyzed in the 20 mM KPO_4_ pH 7.4, 200 mM NaCl with 1% D_2_O.

### Crystallization and data processing

Crystallization was carried out in a sitting drop with 3 mg/ml of the SeMet labeled *V. vulnificus* intra-chain disulfide bond ToxR periplasmic domain (Vv-dbToxRp) mixed 1:1 with 1.7 M ammonium sulfate, .1 M HEPES pH 7.5, and .1 M ammonium formate. The well solution with 40% glucose was used as the cryo-protectant. To cryo-protect the crystals the mother liquor was exchanged with the cryo-protectant in the well. Diffraction data was collected at the NSLS2 FMX beam line set at the Se adsorption edge for SAD data collection. The diffraction data was processed in XDS (Kabsch, 2010) with a space group of P 2_1_ 2_1_ 2_1_ and unit cell of 39.960 40.530 49.570 90.00 90.00 90.00. The structure was solved using PHENIX Auto-Sol and refined using PHENIX Refine (Adams *et al.*, 2010) with COOT (Emsley and Cowtan, 2004) for manual model building. Chimera was used for structure visualization and analysis (Pettersen *et al.*, 2004).

Native Vv-dbToxRp was crystallized by adding in a 1:1 ratio 4 mg/ml of Vv-dbToxRp to 2.0 M ammonium sulfate, and .1 M sodium cacodylate pH 6.3 in a sitting drop. Crystals were cryo-protected by dragging the crystal through 75% paratone N and 25% paraffin oil. Data was collected at NSLS2 FMX beam line. The data was processed using XDS (Kabsch, 2010) with the space group P 2_1_ 2_1_ 2_1_ with a of unit cell of 39.99 40.46 50.29 90.00 90.00 90.00. The structure was solved by molecular replacement using PHASER (McCoy *et al.*, 2007) as implemented in PHENIX (Adams *et al.*, 2010), with the previously solved SeMet structure as the search model. The structure was refined using Refine in PHENIX (Adams *et al.*, 2010) with manual model building in COOT (Emsley and Cowtan, 2004). Chimera was used for structure visualization and analysis (Pettersen *et al.*, 2004).

## Acknowledgments

Funding was provided by NIAID R21-AI140740 and from the BioMt COBRE P20-GM113132. Sequencing was performed by the Molecular Biology Shared Resource at Dartmouth. We thank Dr. Karen Skorupski for her thoughtful comments about the manuscript. We also acknowledge the beam line staff at NSLS2-FMX; Martin Fuchs, Babak Andi, and Wuxian Shi for helping with data collection.

## Authors contributions

FJK and CRM conceived the project and wrote the paper. CRM also performed some of the experiments and solved the X-ray structures. RAS performed experiments as directed, primarily purifying and crystallizing the periplasmic domain for X-ray crystallography. MP performed the NMR experiments, wrote the methods for the NMR experiments, and prepared the HSQC overlay for inclusion in the figures.

## Additional Information

The authors declare no competing interest. All strains, plasmids, and protocols are available upon request. The coordinates for both structures have been deposited in the PDB: SeMet-Vv-dbToxRp, 6uue; native Vv-dbToxRp, 6utc.

## References

Adams, P.D., Afonine, P.V., Bunkoczi, G., Chen, V.B., Davis, I.W., Echols, N., et al. (2010) PHENIX: a comprehensive Python-based system for macromolecular structure solution. Acta Crystallogr D Biol Crystallogr 66: 1–9.

Almagro-Moreno, S., Root, M.Z., and Taylor, R.K. (2015) Role of ToxS in the proteolytic cascade of virulence regulator ToxR in *Vibrio cholerae*. Molecular Microbiology 98: 963–976.

Ante, V.M., Bina, X.R., Howard, M.F., Sayeed, S., Taylor, D.L., and Bina, J.E. (2015) *Vibrio cholerae leuO* transcription is positively regulated by ToxR and contributes to bile resistance. Journal of Bacteriology 197: 3499–3510.

Bina, J., Zhu, J., Dziejman, M., Faruque, S., Calderwood, S., and Mekalanos, J. (2003) ToxR regulon of *Vibrio cholerae* and its expression in vibrios shed by cholera patients. Proc Natl Acad Sci USA 100: 2801–2806.

Champion, G.A., Neely, M.N., Brennan, M.A., and DiRita, V.J. (1997) A branch in the ToxR regulatory cascade of *Vibrio cholerae* revealed by characterization of *toxT* mutant strains. Molecular Microbiology 23: 323–331.

Crawford, J.A., Kaper, J.B., and DiRita, V.J. (1998) Analysis of ToxR-dependent transcription activation of *ompU*, the gene encoding a major envelope protein in *Vibrio cholerae*. Molecular Microbiology 29: 235–246.

DiRita, V.J., and Mekalanos, J.J. (1991) Periplasmic interaction between two membrane regulatory proteins, ToxR and ToxS, results in signal transduction and transcriptional activation. Cell 64: 29–37.

Dziejman, M., Kolmar, H., Fritz, H.J., and Mekalanos, J.J. (1999) ToxR co-operative interactions are not modulated by environmental conditions or periplasmic domain conformation. Molecular Microbiology 31: 305–317.

Emsley, P., and Cowtan, K. (2004) Coot: model-building tools for molecular graphics. Acta Crystallogr D Biol Crystallogr 60: 2126–2132.

Fan, F., Liu, Z., Jabeen, N., Birdwell, L.D., Zhu, J., and Kan, B. (2014) Enhanced interaction of *Vibrio cholerae* virulence regulators TcpP and ToxR under oxygen-limiting conditions. Infection and Immunity 82: 1676–1682.

Fengler, V.H.I., Boritsch, E.C., Tutz, S., Seper, A., Ebner, H., Roier, S., et al. (2012) Disulfide Bond Formation and ToxR Activity in *Vibrio cholerae*. PLoS ONE 7: e47756.

Hase, C.C., and Mekalanos, J.J. (1998) TcpP protein is a positive regulator of virulence gene expression in *Vibrio cholerae*. Proc Natl Acad Sci USA 95: 730–734.

Herrington, D.A., Hall, R.H., Losonsky, G., Mekalanos, J.J., Taylor, R.K., and Levine, M.M. (1988) Toxin, toxin-coregulated pili, and the *toxR* regulon are essential for *Vibrio cholerae* pathogenesis in humans. J Exp Med 168: 1487–1492.

Holm, L., and Laakso, L.M. (2016) Dali server update. Nucleic Acids Res 44: W351–W355.

Hubbard, T.P., Chao, M.C., Abel, S., Blondel, C.J., Abel zur Wiesch, P., Zhou, X., et al. (2016) Genetic analysis of *Vibrio parahaemolyticus* intestinal colonization. Proc Natl Acad Sci USA 113: 6283–6288.

Kabsch, W. (2010) XDS. Acta Crystallogr D Biol Crystallogr 66: 1–8.

Kodama, T., Gotoh, K., Hiyoshi, H., Morita, M., Izutsu, K., Akeda, Y., et al. (2010) Two regulators of *Vibrio parahaemolyticus* play important roles in enterotoxicity by controlling the expression of genes in the Vp-PAI region. PLoS ONE 5: e8678–12.

Kolmar, H., Hennecke, F., Götze, K., Janzer, B., Vogt, B., Mayer, F., and Fritz, H.J. (1995) Membrane insertion of the bacterial signal transduction protein ToxR and requirements of transcription activation studied by modular replacement of different protein substructures. EMBO J 14: 3895–3904.

Krissinel, E., and Henrick, K. (2007) Inference of macromolecular assemblies from crystalline state. J Mol Biol 372: 774–797.

Krukonis, E.S., and DiRita, V.J. (2003) DNA binding and ToxR responsiveness by the wing domain of TcpP, an activator of virulence gene expression in *Vibrio cholerae*. Mol Cell 12: 157–165.

Krukonis, E.S., Yu, R.R., and DiRita, V.J. (2000) The *Vibrio cholerae* ToxR/TcpP/ToxT virulence cascade: distinct roles for two membrane-localized transcriptional activators on a single promoter. Molecular Microbiology 38: 67–84.

Landeta, C., Boyd, D., and Beckwith, J. (2018) Disulfide bond formation in prokaryotes. Nature Microbiology 1–11.

Lembke, M., Pennetzdorfer, N., Tutz, S., Koller, M., Vorkapic, D., Zhu, J., et al. (2018) Proteolysis of ToxR is controlled by cysteine-thiol redox state and bile salts in *Vibrio cholerae*. Molecular Microbiology 1–36.

Li, C.C., Crawford, J.A., DiRita, V.J., and Kaper, J.B. (2000) Molecular cloning and transcriptional regulation of *ompT*, a ToxR-repressed gene in *Vibrio cholerae*. Molecular Microbiology 35: 189–203.

Li, P., Rivera-Cancel, G., Kinch, L.N., Salomon, D., Tomchick, D.R., Grishin, N.V., and Orth, K. (2016) Bile salt receptor complex activates a pathogenic type III secretion system. eLife 5: 1153.

McCoy, A.J., Grosse-Kunstleve, R.W., Adams, P.D., Winn, M.D., Storoni, L.C., and Read, R.J. (2007) Phaser crystallographic software. J Appl Cryst 40: 1–17.

Mey, A.R., Butz, H.A., and Payne, S.M. (2015) *Vibrio cholerae* CsrA regulates ToxR levels in response to amino acids and is essential for virulence. mBio 6: e01064–15–11.

Mey, A.R., Craig, S.A., and Payne, S.M. (2012) Effects of amino acid supplementation on porin expression and ToxR levels in *Vibrio cholerae*. Infection and Immunity 80: 518–528.

Midgett, C.R., Almagro-Moreno, S., Pellegrini, M., Taylor, R.K., Skorupski, K., and Kull, F.J. (2017) Bile salts and alkaline pH reciprocally modulate the interaction between the periplasmic domains of *Vibrio cholerae* ToxR and ToxS. Molecular Microbiology 105: 258–272.

Miller, V.L., and Mekalanos, J.J. (1984) Synthesis of cholera toxin is positively regulated at the transcriptional level by *toxR*. Proceedings of the National Academy of Sciences 81: 3471–3475.

Miller, V.L., and Mekalanos, J.J. (1988) A novel suicide vector and its use in construction of insertion mutations: osmoregulation of outer membrane proteins and virulence determinants in *Vibrio cholerae* requires *toxR*. Journal of Bacteriology 170: 2575–2583.

Miller, V.L., DiRita, V.J., and Mekalanos, J.J. (1989) Identification of *toxS*, a regulatory gene whose product enhances *toxR*-mediated activation of the cholera toxin promoter. Journal of Bacteriology 171: 1288–1293.

Okada, R., Matsuda, S., and Iida, T. (2017) *Vibrio parahaemolyticus* VtrA is a membrane-bound regulator and is activated via oligomerization. PLoS ONE 12: e0187846–16.

Ottemann, K.M., and Mekalanos, J.J. (1995) Analysis of *Vibrio cholerae* ToxR function by construction of novel fusion proteins. Molecular Microbiology 15: 719–731.

Ottemann, K.M., and Mekalanos, J.J. (1996) The ToxR protein of *Vibrio cholerae* forms homodimers and heterodimers. Journal of Bacteriology 178: 156–162.

Pettersen, E.F., Goddard, T.D., Huang, C.C., Couch, G.S., Greenblatt, D.M., Meng, E.C., and Ferrin, T.E. (2004) UCSF Chimera--a visualization system for exploratory research and analysis. J Comp Biol 25: 1605–1612.

Provenzano, D., Schuhmacher, D.A., Barker, J.L., and Klose, K.E. (2000) The virulence regulatory protein ToxR mediates enhanced bile resistance in *Vibrio cholerae* and other pathogenic *Vibrio* species. Infection and Immunity 68: 1491–1497.

Reich, K.A., and Schoolnik, G.K. (1994) The light organ symbiont *Vibrio fischeri* possesses a homolog of the *Vibrio cholerae* transmembrane transcriptional activator ToxR. Journal of Bacteriology.

Schleucher, J., Schwendinger, M., Sattler, M., Schmidt, P., Schedletzky, O., Glaser, S.J., et al. (1994) A general enhancement scheme in heteronuclear multidimensional NMR employing pulsed field gradients. J Biomol NMR 4: 301–306.

Schlundt, A., Buchner, S., Janowski, R., Heydenreich, T., Heermann, R., Lassak, J.X.R., et al. (2017) Structure-function analysis of the DNA-binding domain of a transmembrane transcriptional activator. Nature Publishing Group 1–16.

Sivashanmugam, A., Murray, V., Cui, C., Zhang, Y., Wang, J., and Li, Q. (2009) Practical protocols for production of very high yields of recombinant proteins using *Escherichia coli*. Protein Sci 18: 936–948.

Studier, F.W. (2005) Protein production by auto-induction in high-density shaking cultures. Protein Expr Purif 41: 207–234.

Wang, S.-Y., Lauritz, J., Jass, J., and Milton, D.L. (2002) A ToxR homolog from *Vibrio anguillarum* serotype O1 regulates its own production, bile resistance, and biofilm formation. Journal of Bacteriology 184: 1630–1639.

Welch, T.J., and Bartlett, D.H. (1998) Identification of a regulatory protein required for pressure-responsive gene expression in the deep-sea bacterium *Photobacterium species strain SS9*. Molecular Microbiology 27: 977–985.

Xue, Y., Tu, F., Shi, M., Wu, C.-Q., Ren, G., Wang, X., et al. (2016) Redox Pathway Sensing Bile Salts Activates Virulence Gene Expression in *Vibrio cholerae*. Molecular Microbiology 1–16.

Yang, Y., and Isberg, R.R. (1997) Transcriptional regulation of the *Yersinia pseudotuberculosis* pH 6 antigen adhesin by two envelope-associated components. Molecular Microbiology 24: 499–510.

