## Supplemental information for "A disulfide constrains the ToxR periplasmic domain structure, altering its interactions with ToxS and bile-salts"

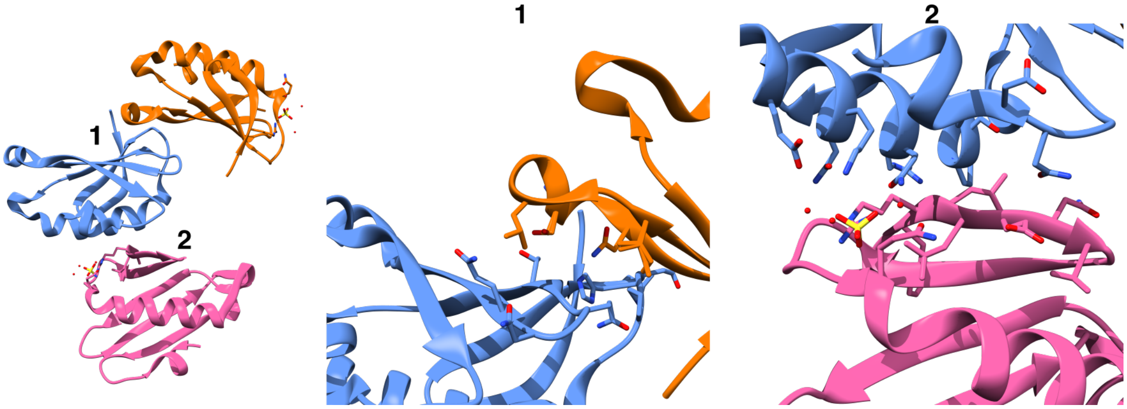


**Figure S1:** Figure showing crystal contacts that were explored as potential dimerization interfaces. Left panel shows how the monomers contact each other in the crystal. Detail of the first (middle panel) and the second (right panel) interface.

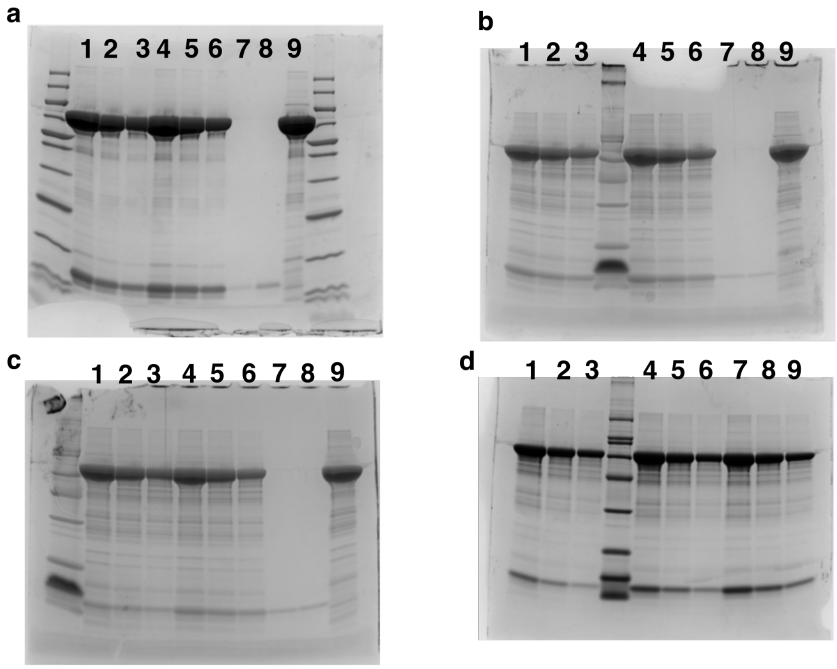


**Figure S2:** Gels of the pull downs excerpted in figure 1 and 2. **A** is the gel from the validation that the dbToxRp is pulled down by ToxS. Lanes 1-3 are the 1x, 1/2x, and 1/4x dilutions of the non-disulfide containing ToxR periplasmic domain. Lanes 4-6 are the dilution series of the dbToxRp pull down. Lane 7 is the dbToxRp only control, lane 8 is the ToxRp only control, and lane 9 is the ToxSp lysate only control. **B** is the gel of the dbToxRp pull down with and without CDC. Lanes 1-3 are the 1x, 1/2x, 1/4x dilution series of dbToxRp without CDC and lanes 4-6 are the dilution series of dbToxRp with CDC. Lane 8 is the dbToxRp only control without CDC, lane 9 is the dbToxRp control with CDC and lane 10 is CBDI-ToxSp only control. **C** is the ToxRp pull down with and without CDC, lanes 1-3 is the buffer treated dilution series, lanes 4-6 are the CDC treated dilution series, lane 7 is the ToxRp only control, lane 8 is the ToxRp with CDC control, and lane 9 is the CBDI-ToxSp only control. **D** is the gel comparing the amount of ToxRp pulled down relative to dbToxRp. Lanes 1-3 is the dilution series of ToxRp with buffer, and lanes 4-6 is the dilution series of ToxRp with CDC, and lanes 7-9 is the dilution series of dbToxRp with CDC.

Table S1: DALI table excerpt

| # No | Chain | rmsd | %id | Description |
|---|---|---|---|---|
| 118 | 5kew-E | 3 | 8 | VTRA PROTEIN; |
| 119 | 6roa-B | 3.2 | 8 | CYSTATIN-C; |
| 120 | 1j0x-P | 2 | 10 | GLYCERALDEHYDE-3-PHOSPHATE DEHYDROGENASE; |
| 121 | 3ub1-A | 4.2 | 4 | ORF13-LIKE PROTEIN; |
| 122 | 5kew-C | 3.2 | 8 | VTRA PROTEIN; |
| 123 | 1pp6-D | 2.5 | 9 | VOLVATOXIN A2; |
| 124 | 6gve-I | 1.8 | 8 | CP12 POLYPEPTIDE; |
| 125 | 3nx0-A | 3.3 | 8 | CYSTATIN-C; |
| 126 | 3wz4-H | 3 | 11 | DOTI; |
| 127 | 2xh9-B | 3.6 | 4 | ORF12; |
| 128 | 2vt8-B | 2.3 | 13 | PROTEASOME INHIBITOR PI31 SUBUNIT; |
| 129 | 6ct5-B | 3.5 | 6 | 4'-PHOSPHOPANTETHEINYL TRANSFERASE; |
| 130 | 2bo9-B | 3.6 | 8 | CARBOXYPEPTIDASE A4; |
| 131 | 3ezy-B | 2.3 | 7 | DEHYDROGENASE; |
| 132 | 5kew-A | 3.5 | 6 | VTRA PROTEIN; |

**Table S1**: Table showing excerpted DALI results of the dbToxRp structure. Highlighted in yellow are the alignments with the VtrA structure.

Table S2 Refinement Statistics

| | Native_VvToxRp | SeMet-VvToxRp |
|---|---|---|
| Wavelength (Å) | 0.97933 | 0.97934 |
| Resolution range | 28.43 - 1.249 (1.294 - 1.249) | 24.78 - 1.389 (1.439 - 1.389) |
| Space group | P 21 21 21 | P 21 21 21 |
| Unit cell | 39.97 40.44 50.28 90 90 90 | 39.96 40.53 49.57 90 90 90 |
| Total reflections | 229089 (5713) | 202709 (13693) |
| Unique reflections | 20417 (1084) | 16768 (1604) |
| Multiplicity | 11.2 (5.3) | 12.1 (8.5) |
| Completeness (%) | 87.81 (47.08) | 98.73 (92.82) |
| Anomalous Completeness (%) | | 99.66 |
| Mean I/sigma(I) | 23.88 (1.16) | 21.66 (1.20) |
| Wilson B-factor | 17.28 | 21.45 |
| R-merge | 0.0473 (0.9164) | 0.06322 (1.182) |
| R-meas | 0.04944 (1.016) | 0.06591 (1.258) |
| R-pim | 0.0141 (0.4206) | 0.0184 (0.4185) |
| CC1/2 | 1 (0.446) | 0.999 (0.636) |
| CC* | 1 (0.786) | 1 (0.882) |
| Reflections used in refinement | 20408 (1081) | 16585 (1512) |
| Reflections used for R-free | 1980 (108) | 1654 (149) |
| R-work | 0.1955 (0.2853) | 0.2007 (0.2863) |
| R-free | 0.2118 (0.2857) | 0.2287 (0.3124) |
| CC(work) | 0.964 (0.721) | 0.965 (0.819) |
| CC(free) | 0.955 (0.684) | 0.963 (0.809) |
| Number of non-hydrogen atoms | 848 | 770 |
| macromolecules | 737 | 708 |
| ligands | 5 | 5 |
| solvent | 106 | 57 |
| Protein residues | 91 | 91 |
| RMS(bonds) | 0.006 | 0.006 |
| RMS(angles) | 0.86 | 0.81 |
| Ramachandran favored (%) | 96.63 | 98.88 |
| Ramachandran allowed (%) | 3.37 | 1.12 |
| Ramachandran outliers (%) | 0 | 0 |
| Rotamer outliers (%) | 2.35 | 0 |
| Clashscore | 4.73 | 2.82 |
| Average B-factor | 23.7 | 27.72 |
| macromolecules | 22.44 | 27.05 |
| ligands | 23.34 | 27.76 |
| solvent | 32.5 | 35.96 |
